# A chromosome-level genome assembly of northern dragonhead, *Dracocephalum ruyschiana* L., 1753

**DOI:** 10.64898/2026.11.11.765931

**Authors:** Marius A. Strand, Marianne Evju, Magni Olsen Kyrkjeeide, Ole K. Tørresen, Bram Danneels, Mariann Árnyasi, Giada Ferrari, Morten Skage, Ave Tooming-Klunderud, Kjetill S. Jakobsen

**Affiliations:** Department of Biosciences, University of Oslo, Oslo, Norway; Norwegian Institute for Nature Research (NINA), Oslo, Norway; Norwegian Institute for Nature Research (NINA), Trondheim, Norway; Computational Biology Unit (CBU), Department of Informatics, University of Bergen, Bergen, Norway; Department of Animal and Aquacultural Sciences, Faculty of Biosciences, Norwegian University of Life Sciences (NMBU), Ås, Norway

## Abstract

We present a chromosome-level genome assembly for the vascular plant northern dragonhead, *Dracocephalum ruyschiana*. The assembly comprises two pseudo-haplotypes, hap1 and hap2, of 661 Mb and 681 Mb, with 93.6% and 92.5%, respectively, scaffolded into seven chromosomes each – in line with the known karyotype. Assembly quality was high, with conserved single-copy ortholog completeness scores of 97.9% and 97.8%, combined k-mer completeness of 98.1%, and 38,141 and 38,617 annotated protein-coding genes for hap1 and hap2, respectively. This assembly provides a high-quality genomic resource for future research and conservation of northern dragonhead, as well as the endangered northern dragonhead sap beetle, which depends on the plant for reproduction and whose genome assembly is described in a complementary genome note.

## Introduction

Northern dragonhead (*Dracocephalum ruyschiana*) is a perennial, long-lived plant in the Lamiaceae family (Lid & Lid, 2005; Milberg & Bertilsson, 1997). It has a Eurasian distribution, from Norway in the northwest to the Pyrenees in the southwest, scattered through Europe and Russia to Mongolia and Japan (Bánki et al., 2026). The genus *Dracocephalum* is found in the northern temperate zone, and includes about 70 species, with a distribution mainly confined to dry steppe in Central and West Asia, as well as in the alpine region of the Qinghai-Tibetan Plateau (Chen et al., 2022). Species within the genus are widely known for their pharmacological and medicinal properties (Liu et al., 2024). Northern dragonhead is one of few species in the genus occurring in Europe.

Northern dragonhead usually grows on relatively dry, open, often limestone-dominated habitats, including mountain slopes, dry grasslands, open forests, and rocky outcrops (Lazarević et al., 2009). In Northern Europe, the species is found mainly below the forest line, on calcareous soils, whereas in Central Europe, northern dragonhead is found in alpine and subalpine areas, on dry mountain hills. The Russian populations are associated with the large steppe habitats. The occurrences are highly fragmented, and the species’ distribution suggests that current populations constitute relicts from a more continuous distribution. Northern dragonhead likely survived the Quaternary ice ages in ice-free areas of continental Asia, and as the ice retreated at the end of the last glacial period, the species gained the opportunity to spread westward through the steppe-like vegetation that for a time dominated large parts of Europe (Hegi, 1964; Lazarević et al., 2009). Subsequently, forests came to dominate the lowlands, causing many northern dragonhead populations to be shaded out (Blytt, 1876; Gjærevoll, 1992; Sterner, 1922). As agriculture and livestock farming gradually shaped the European landscape through cultivation and grazing, suitable lowland habitats for the northern dragonhead once again became available.

Northern dragonhead is diploid (2n = 14) (Lid & Lid, 2005). Plants normally reach about 30 cm. Multiple stems arise from a rhizome, which can be large, and highly branched (Lid & Lid, 2005), suggesting that genets can reach a significant age. The lateral vegetative spread is, nevertheless, assumed to be limited (Klimešová et al., 2017).

The stems bear large, blue flowers, 2–2.5 cm long, that are assembled in spike-like inflorescences (Figure 1B). Northern dragonhead typically flowers in June, and bumblebees are the most important pollinators, although the species is self-compatible (Milberg & Bertilsson, 1997). Each flower produces up to four seeds, which are relatively large (2 −3 mm) without obvious adaptations for long-distance dispersal (Milberg & Bertilsson, 1997). Limited dispersibility is confirmed by genetic data showing that northern dragonhead is genetically structured at a regional scale (Kyrkjeeide et al., 2020).

**Figure 1.**
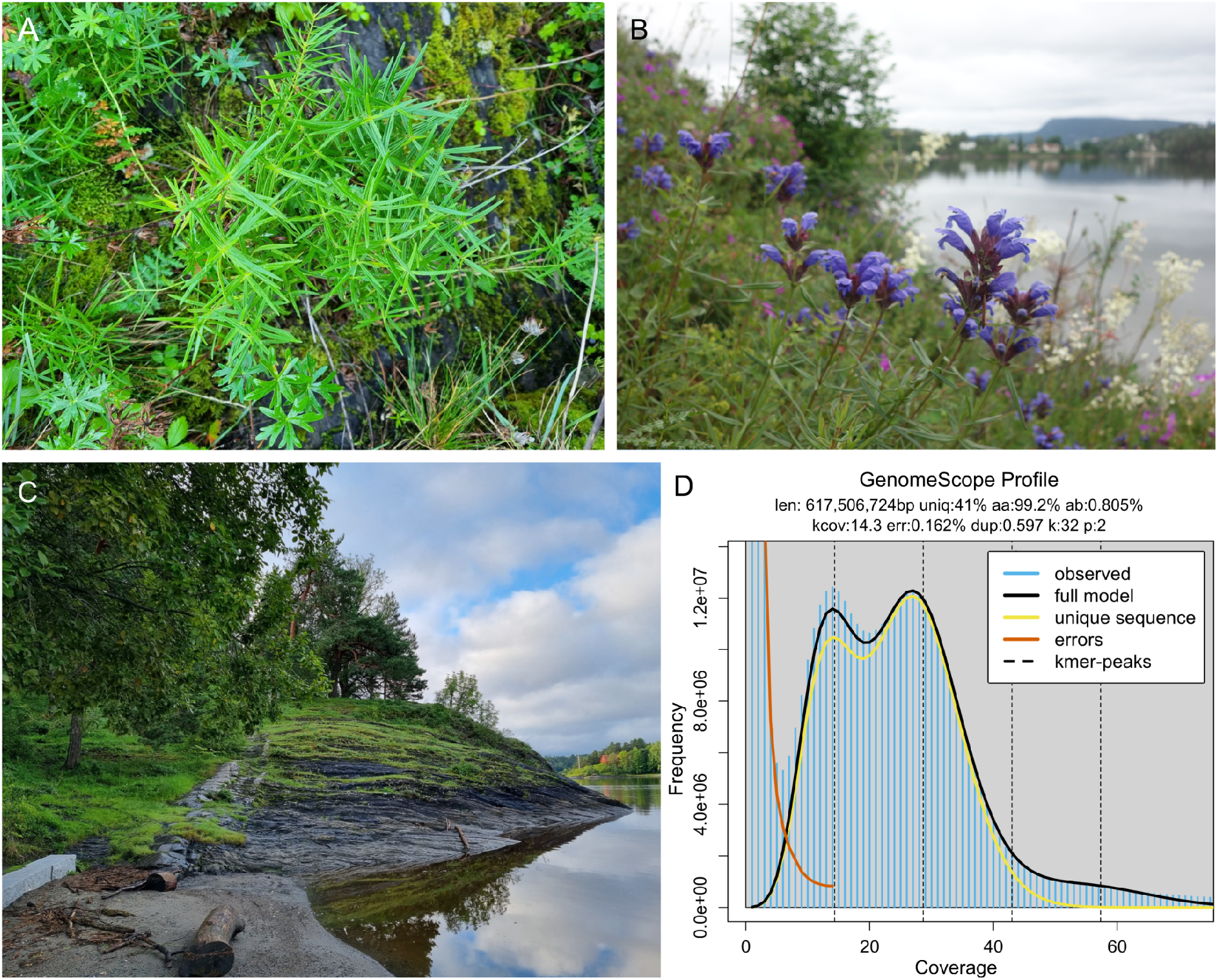
Sequenced specimen and genome profile. **A)** Sequenced individual of northern dragonhead, *Dracocephalum ruyschiana*, photo by Marianne Evju. **B)** Northern dragonhead in bloom; photo by Marianne Evju. **C)** Collection environment of the sequenced northern dragonhead specimen; photo by Marianne Evju. **D)** GenomeScope profile of the HiFi reads from the sequenced individual. This analysis estimates a 617.5 Mb genome, with 0.805% heterozygosity. The left-hand peak of k-mers corresponds to k-mers from heterozygous regions of the genome, while the right-hand peak is from homozygous regions.

On the European Red List, the northern dragonhead is considered Least Concern (Ericsson et al., 2011), due to its apparently stable populations in Russia. In Europe, however, the species is listed under Appendix I of the Bern Convention (1979), and it is considered threatened in several national and regional Red Lists (Ginszt & Laskowska-Ginszt, 2022; Reteau, 2019; Sundberg et al., 2020; Tomović et al., 2009), including Norway, where it is listed as Vulnerable (Solstad et al., 2021), due to an estimated 20 −40% decline over the last 30 years. Changes in land use, particularly related to changing agricultural practices, with reduced use of extensive management and increased intensification, are the main causes of decline, in addition to land conversion due to development projects (housing, infrastructure, etc.).

Northern dragonhead is also the host plant of the endangered northern dragonhead sap beetle (*Thymogethes norvegicus*) which is believed to be endemic to Norway (Artsdatabanken, 2021) and depends on the plant for reproduction (Stabbetorp & Endrestøl, 2011). The genome of the northern dragonhead sap beetle is presented in a companion article (Strand et al., 2026).

Northern dragonhead reaches its northwestern distribution limit in Norway (Bánki et al., 2026), where currently >25% of the European (excluding Russia) populations can be found. Thus, conserving it in Norway is of importance to maintain the European population. In Norway, its distribution is southeastern, in areas with warm summers, reaching an altitudinal limit of approximately 880 m asl.

Here, we present a chromosome-level, haplotype-resolved genome assembly of the northern dragonhead generated using PacBio HiFi in combination with Hi-C data. The public availability of this genome resource will support future research and conservation efforts of northern dragonhead. This genome assembly was generated as part of the Earth BioGenome Project Norway.

## Material and Methods

### Sample acquisition and DNA extraction

Five shoots from a non-flowering ramet of northern dragonhead were collected in Bærum municipality, Akershus, Norway (59.88412° N, 10.53523° E), on 28 August 2023 (Figure 1A,C). The ramet consisted of 19 shoots, of which five were sampled using tweezers to detach the shoots. A permit for sampling was provided by the County Governor oføstfold, Buskerud, Oslo og Akershus (permit no. 2023/29186), dated 25 August 2023. The sample (daDraRuys1) was stored at −80 °C.

### Library preparation and sequencing for *de novo* assembly

A total of 720 mg of leaf tissue was ground to a fine powder using a mortar and pestle under liquid nitrogen. High-molecular-weight (HMW) DNA was isolated using the NucleoBond HMW DNA kit (Macherey-Nagel), following the manufacturer’s protocol. Quality check of the amount, purity and integrity of the isolated DNA was performed using a combination of Qubit BR DNA quantification assay kit (Thermo Fisher), Nanodrop (Thermo Fisher), and Fragment Analyser (DNA HS 50kb large fragment kit, Agilent Tech.).

Before PacBio HiFi library preparation, DNA was purified an additional time using AMPure PB beads (1:1 ratio). Purified HMW DNA was sheared to an average fragment size of 15–20 kb large fragments using the Megaruptor3 (Diagenode). A HiFi library was prepared following the PacBio protocol for HiFi library preparation using the SMRTbell® Prep Kit 3.0. The final HiFi library was size-selected with a 10 kb cut-off using a BluePippin (Sage Sciences) and sequencing was performed by the Norwegian Sequencing Centre on a PacBio Revio instrument (Pacific Biosciences Inc). The library was sequenced as part of a pool over three 25M SMRT cells with Revio sequencing chemistry.

A Hi-C library was prepared using the Arima High Coverage HiC kit (Arima Genomics), following the manufacturer’s recommendations and starting with 1300 mg of leaf tissue. Chromatin was crosslinked prior to nuclei isolation as written in the user guide for plant tissues (Doc A160163 v01). Final library quality was quantified using a Kapa Library quantification kit for Illumina (Roche Inc.). The library was sequenced with other libraries on an Illumina NovaSeq X (Illumina Inc) with 2*150 bp paired end mode on a 25B flowcell at the Norwegian Sequencing Centre (https://www.sequencing.uio.no).

### Genome assembly and curation, annotation and evaluation

A full list of relevant software tools and versions is presented in Table 1. We assembled the species using a pre-release of the Earth BioGenome Project Norway (EBP-Nor) genome assembly pipeline (https://github.com/ebp-nor/GenomeAssembly). The main steps of this pipeline are described below. KMC (Kokot et al., 2017) was used to count k-mers of size 32 in the PacBio HiFi reads, excluding k-mers occurring more than 10,000 times. GenomeScope (Ranallo-Benavidez et al., 2020) was run as part of the pipeline on the k-mer histogram output from KMC to estimate genome size, heterozygosity and repeat content (Figure 1D). Ploidy level was investigated using Smudgeplot (Ranallo-Benavidez et al., 2020). HiFiAdapterFilt (Sim et al., 2022) was applied on the HiFi reads to remove possible remnant PacBio adapter sequences. The filtered HiFi reads were assembled using hifiasm (Cheng et al., 2021) with Hi-C integration resulting in a pair of haplotype-resolved assemblies, pseudo-haplotype one (hap1) and pseudo-haplotype two (hap2). Unique k-mers in each assembly/pseudo-haplotype were identified using meryl (Rhie et al., 2020) and used to create two sets of Hi-C reads, one without any k-mers occurring uniquely in hap1 and the other without k-mers occurring uniquely in hap2.

**Table 1.** Software tools: versions and sources.

| Software tool | Version | Source |
|---|---|---|
| AGAT | 1.4 | https://github.com/NBISweden/AGAT |
| BEDTools | 2.31.1 | https://github.com/arq5x/bedtools2 |
| blobtk | 0.5.8 | https://github.com/blobtoolkit/blobtk |
| BlobToolKit | 4.1.7 | https://github.com/blobtoolkit/blobtoolkit |
| BUSCO | 5.8.3 | https://gitlab.com/ezlab/busco |
| BWA-MEM | 0.7.18 | https://github.com/lh3/bwa |
| DIAMOND | 2.1.8 | https://github.com/bbuchfink/diamond |
| EMBLmyGFF3 | 2.2 | https://github.com/NBISweden/EMBLmyGFF3 |
| EvidenceModeler | 2.1.0 | https://github.com/EVidenceModeler/EVidenceModeler |
| FCS-GX | 0.4.0 | https://github.com/ncbi/fcs |
| Funannotate | 1.8.17 | https://github.com/nextgenusfs/funannotate |
| GALBA | 1.0.9 | https://github.com/Gaius-Augustus/GALBA |
| GenomeScope | 2.0 | https://github.com/tbenavi1/genomescope2.0 |
| Gfastats | 1.3.6 | https://github.com/vgl-hub/gfastats |
| Helixer | 0.3.5 | https://github.com/usadellab/Helixer |
| HiFiAdapterFilt | 2.0.0 | https://github.com/sheinasim/HiFiAdapterFilt |
| hifiasm | 0.20.0 | https://github.com/chhylp123/hifiasm |
| InterProScan | 5.47-82 | https://www.ebi.ac.uk/interpro/search/sequence/ |
| KMC | 3.1.1 | https://github.com/refresh-bio/KMC |
| Merqury | 1.3 | https://github.com/marbl/merqury |
| meryl | 1.3.0 | https://github.com/marbl/meryl |
| minimap2 | 2.28 | https://github.com/lh3/minimap2 |
| miniprot | 0.11-r234 | https://github.com/lh3/miniprot |
| Oatk | 1.0 | https://github.com/c-zhou/oatk |
| MitoHiFi | 3.0.0 | https://github.com/marcelauliano/MitoHiFi |
| PretextMap | 0.1.9 | https://github.com/wtsi-hpag/PretextMap |
| PretextSnapshot | - | https://github.com/wtsi-hpag/PretextSnapshot |
| PretextView | 0.2.5 | https://github.com/wtsi-hpag/PretextView |
| Rapid curation 2.0 | 964d17e997e00c69f25940cf96d3658bda631147 | https://github.com/Nadolina/Rapid-curation-2.0, |
| RED | 2018.09.10 | https://github.com/BioinformaticsToolsmith/Red |
| samtools | 1.17 | https://github.com/samtools/samtools |
| smudgeplot | 0.2.5 | https://github.com/KamilSJaron/smudgeplot |
| YaHS | 1.2.2 | https://github.com/c-zhou/yahs |

K-mer filtered Hi-C reads were aligned to each scaffolded assembly using BWA-MEM (Li, 2013) with -5SPM options. The alignments were sorted based on name using samtools (Li et al., 2009) before applying samtools fixmate to remove unmapped reads and secondary alignments and to add mate score, and samtools markdup to remove duplicates. The resulting BAM files were used to scaffold the two assemblies using YaHS (Zhou et al., 2023) with default options. FCS-GX (Astashyn et al., 2024) was used to search for putative contamination and contaminated sequences were removed. The mitochondria and plastids were searched for in contigs and reads using both Oatk (Zhou et al., 2025) and MitoHifi (Uliano-Silva et al., 2023), identifying the chloroplast genome but not the mitochondrial genome. The assemblies were manually curated using PretextView and Rapid curation 2.0.

We annotated the genome assemblies using a pre-release version of the EBP-Nor genome annotation pipeline (https://github.com/ebp-nor/GenomeAnnotation). The main steps of this pipeline are described below. First, AGAT (https://zenodo.org/record/7255559) agat_sp_keep_longest_isoform.pl and agat_sp_extract_sequences.pl were used on the *Arabidopsis thaliana* (TAIR10.1) genome assembly and annotation to generate one protein (the longest isoform) per gene. Miniprot (Li, 2023) was used to align the proteins to the curated assemblies. UniProtKB/Swiss-Prot (UniProt Consortium, 2023) release 2026_03 in addition to the Viridiplantae part of OrthoDB v12 (Tegenfeldt et al., 2025) were also aligned separately to the assemblies. RED (Girgis, 2015) was run via redmask (https://github.com/nextgenusfs/redmask) on the assemblies to mask repetitive areas. In addition to gene models from protein alignments, GALBA (Brůna et al., 2023; Buchfink et al., 2015; Hoff & Stanke, 2019; Li, 2023; Stanke et al., 2006) was run with the *Arabidopsis thaliana* proteins using the miniprot mode on the masked assemblies to generate *ab initio* predicted gene models. Helixer (Holst et al., 2025) was run using the land plant model (land_plant_v0.3_a_0080.h5). The funannotate-runEVM.py script from Funannotate was used to run EvidenceModeler (Haas et al., 2008) on the alignments of TAIR10.1 proteins, UniProtKB/Swiss-Prot proteins, Viridiplantae proteins and the predicted genes from GALBA and Helixer. The resulting predicted proteins were compared to the protein repeats that Funannotate distributes using DIAMOND blastp and the predicted genes were filtered based on this comparison using AGAT. The filtered proteins were compared to the UniProtKB/Swiss-Prot release 2026_03 using DIAMOND (Buchfink et al., 2015) blastp to find gene names and InterProScan (Jones et al., 2014) was used to discover functional domains. AGATs agat_sp_manage_functional_annotation.pl was used to attach the gene names and functional annotations to the predicted genes. EMBLmyGFF3 (Norling et al., 2018) was used to combine the FASTA files and GFF3 files into a EMBL format for submission to ENA.

All the evaluation tools have also been implemented in a pipeline, similar to assembly and annotation (https://github.com/ebp-nor/GenomeEvaluation). The main steps of this pipeline are described below. Merqury (Rhie et al., 2020) was used to assess the completeness and quality of the genome assemblies by comparing them to the k-mer content of both the Hi-C reads and PacBio HiFi reads. BUSCO (Manni et al., 2021) was used to assess the completeness of the genome assemblies by comparing against the expected gene content in the lamiales (odb12.2) lineage. Gfastats (Formenti et al., 2022) was used to output different assembly statistics of the assemblies. BlobToolKit and BlobTools2 (Laetsch & Blaxter, 2017), in addition to blobtk were used to visualize assembly statistics. To generate the Hi-C contact map image, the Hi-C reads were mapped to the assemblies using BWA-MEM (Li, 2013) using the same approach as above. Finally, PretextMap was used to create a contact map which was visualized using PretextSnapshot.

To characterize the differences between the two pseudo-haplotypes, we ran a genome alignment using minimap2 (Li, 2018) on the homologous chromosomes. The resulting alignment was processed with the minimap2-included paftools.js producing a report listing the number of substitutions, insertions and deletions between the two pseudo-haplotypes.

The text in Methods and parts of Results is based on a template we use for all the species we publish in the EBP-Nor project.

## Results

### *De novo* genome assembly and annotation

The genome of northern dragonhead was assembled from a total of 30-fold coverage in Pacific Biosciences single-molecule HiFi long reads and 57-fold coverage in Arima Hi-C reads resulting in two haplotype-separated assemblies. The final assemblies have total lengths of 661 Mb and 681 Mb (Table 2 and Figure 2), respectively. Pseudo-haplotypes one (hap1) and two (hap2) have scaffold N50 size of 93.4 Mb and 94.9 Mb, respectively, and contig N50 size of 3.9 Mb and 5.1 Mb, respectively (Table 2 and Fig. 2). Seven pseudo-chromosomes were identified in both pseudo-haplotypes. Chromosomes are named according to length in descending order.

**Table 2.**
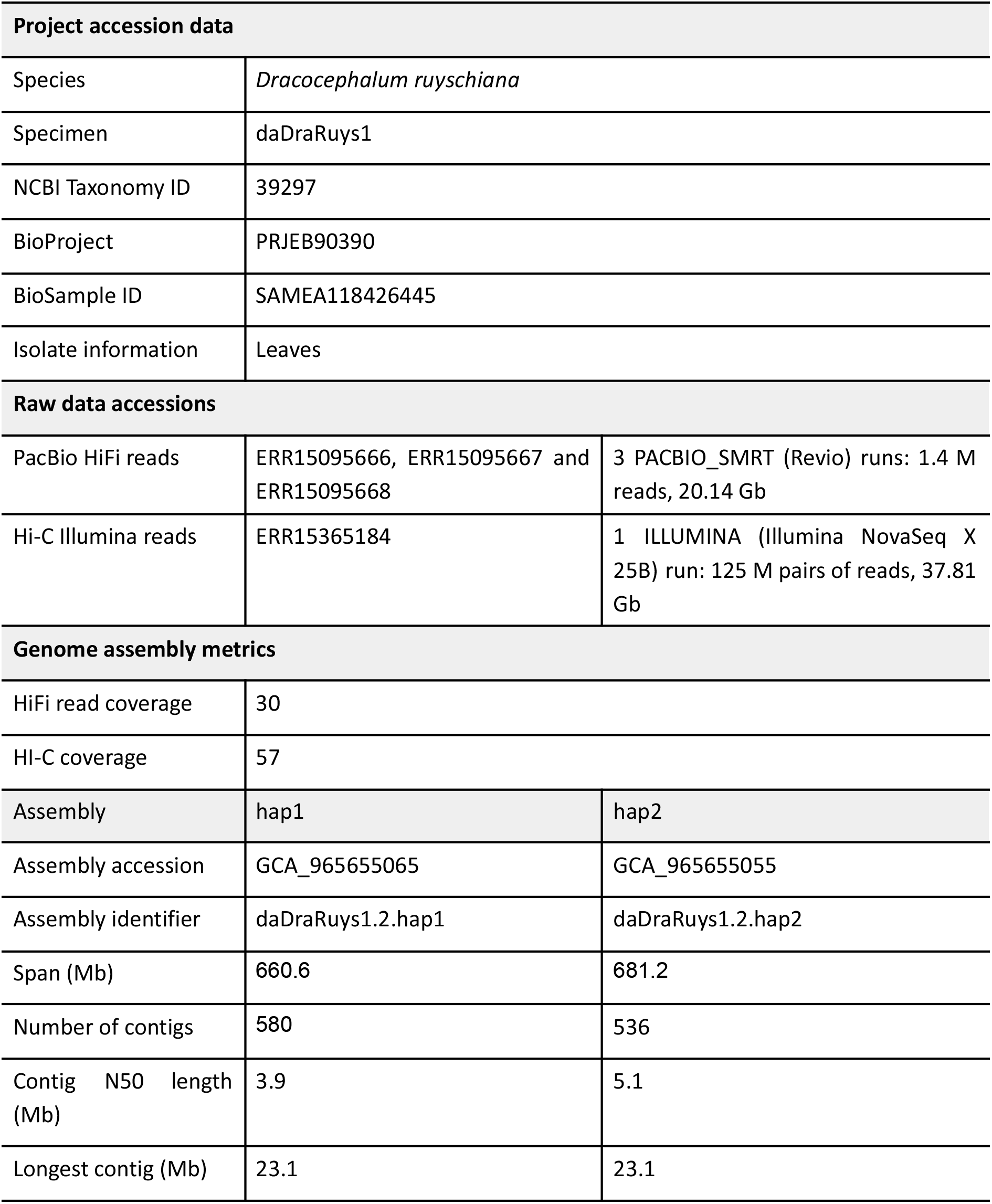
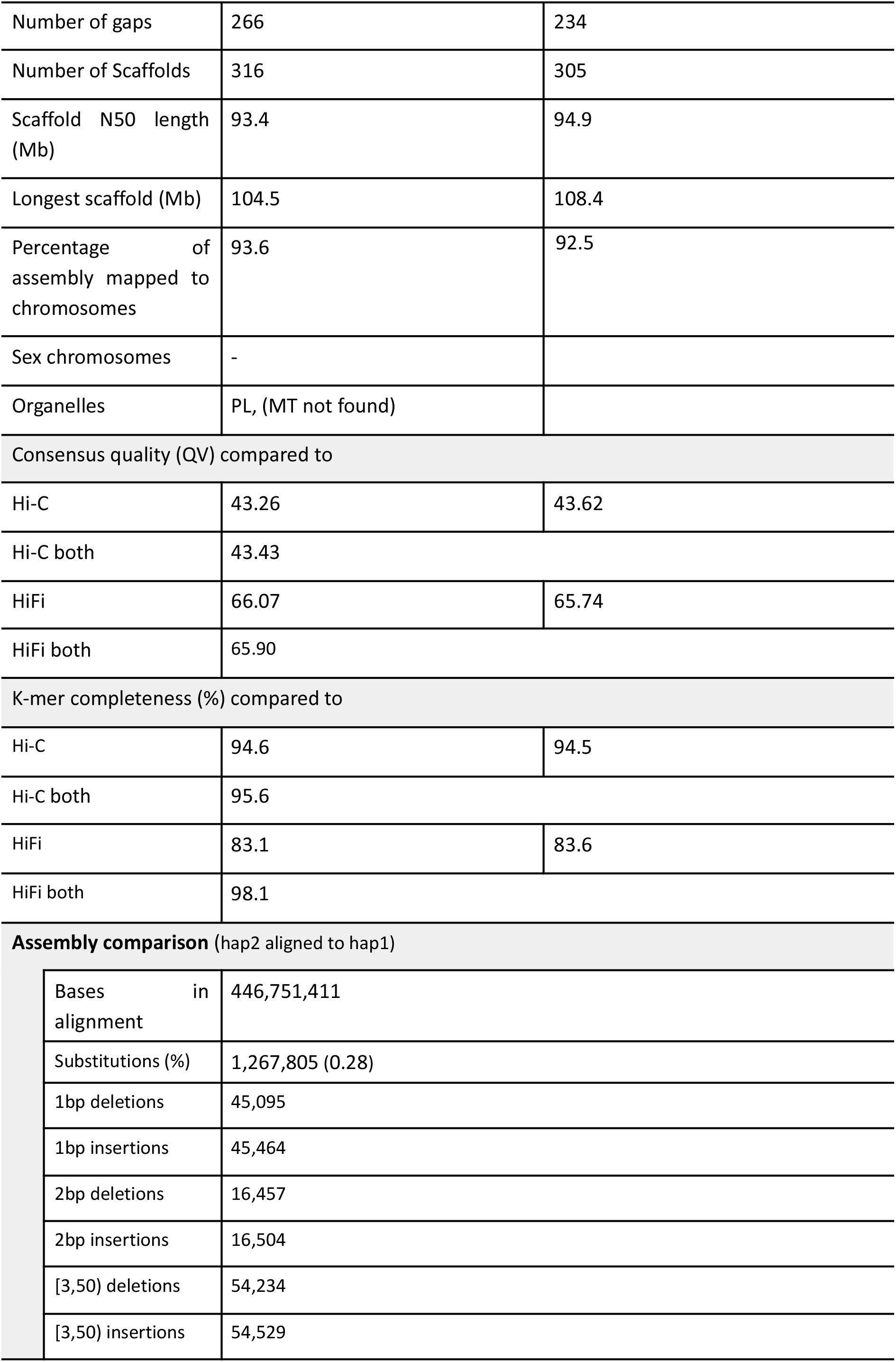
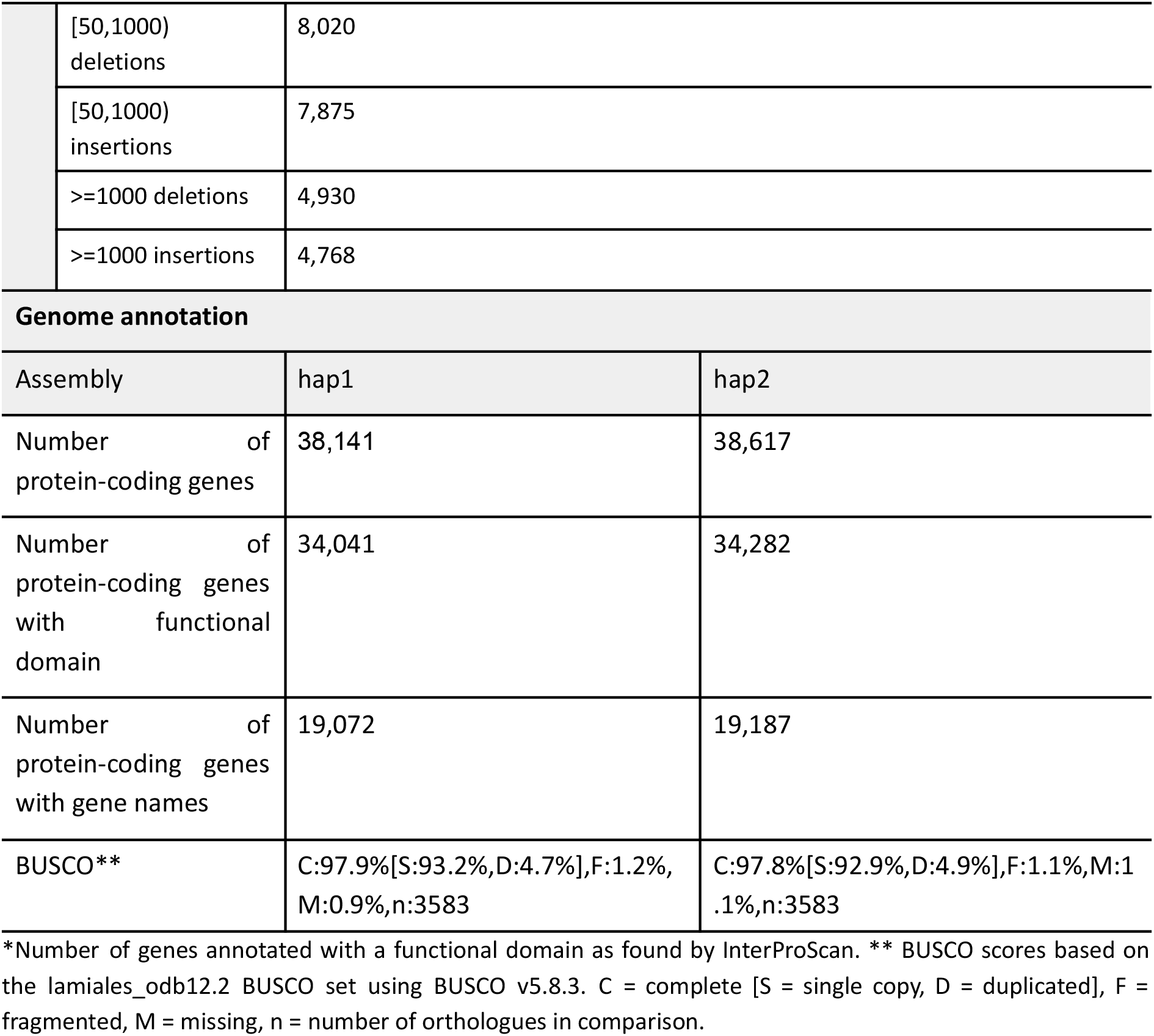
Genome data for *Dracocephalum ruyschiana*.

| **Project accession data** |  |  |
|---|---|---|
| Species | *Dracocephalum ruyschiana* |  |
| Specimen | daDraRuys1 |  |
| NCBI Taxonomy ID | 39297 |  |
| BioProject | PRJEB90390 |  |
| BioSample ID | SAMEA118426445 |  |
| Isolate information | Leaves |  |
| **Raw data accessions** |  |  |
| PacBio HiFi reads | ERR15095666, ERR15095667 and ERR15095668 | 3 PACBIO_SMRT (Revio) runs: 1.4 M reads, 20.14 Gb |
| Hi-C Illumina reads | ERR15365184 | 1 ILLUMINA (Illumina NovaSeq X 25B) run: 125 M pairs of reads, 37.81 Gb |
| **Genome assembly metrics** |  |  |
| HiFi read coverage | 30 |  |
| HI-C coverage | 57 |  |
| Assembly | hap1 | hap2 |
| Assembly accession | GCA_965655065 | GCA_965655055 |
| Assembly identifier | daDraRuys1.2.hap1 | daDraRuys1.2.hap2 |
| Span (Mb) | 660.6 | 681.2 |
| Number of contigs | 580 | 536 |
| Contig N50 length (Mb) | 3.9 | 5.1 |
| Longest contig (Mb) | 23.1 | 23.1 |
| Number of gaps | 266 | 234 |
| Number of Scaffolds | 316 | 305 |
| Scaffold N50 length (Mb) | 93.4 | 94.9 |
| Longest scaffold (Mb) | 104.5 | 108.4 |
| Percentage of assembly mapped to chromosomes | 93.6 | 92.5 |
| Sex chromosomes | - |  |
| Organelles | PL, (MT not found) |  |
| Consensus quality (QV) compared to |  |  |
| Hi-C | 43.26 | 43.62 |
| Hi-C both | 43.43 |  |
| HiFi | 66.07 | 65.74 |
| HiFi both | 65.90 |  |
| K-mer completeness (%) compared to |  |  |
| Hi-C | 94.6 | 94.5 |
| Hi-C both | 95.6 |  |
| HiFi | 83.1 | 83.6 |
| HiFi both | 98.1 |  |
| **Assembly comparison** (hap2 aligned to hap1) |  |  |
| Bases in alignment | 446,751,411 |  |
| Substitutions (%) | 1,267,805 (0.28) |  |
| 1bp deletions | 45,095 |  |
| 1bp insertions | 45,464 |  |
| 2bp deletions | 16,457 |  |
| 2bp insertions | 16,504 |  |
| [3,50) deletions | 54,234 |  |
| [3,50) insertions | 54,529 |  |
| [50,1000) deletions | 8,020 |  |
| [50,1000) insertions | 7,875 |  |
| >=1000 deletions | 4,930 |  |
| >=1000 insertions | 4,768 |  |
| **Genome annotation** |  |  |
| Assembly | hap1 | hap2 |
| Number of protein-coding genes | 38,141 | 38,617 |
| Number of protein-coding genes with functional domain | 34,041 | 34,282 |
| Number of protein-coding genes with gene names | 19,072 | 19,187 |
| BUSCO** | C:97.9%[S:93.2%,D:4.7%],F:1.2%,M:0.9%,n:3583 | C:97.8%[S:92.9%,D:4.9%],F:1.1%,M:1.1%,n:3583 |

\*Number of genes annotated with a functional domain as found by InterProScan. \*\* BUSCO scores based on the lamiales_odb12.2 BUSCO set using BUSCO v5.8.3. C = complete [S = single copy, D = duplicated], F = fragmented, M = missing, n = number of orthologues in comparison.

**Figure 2.**
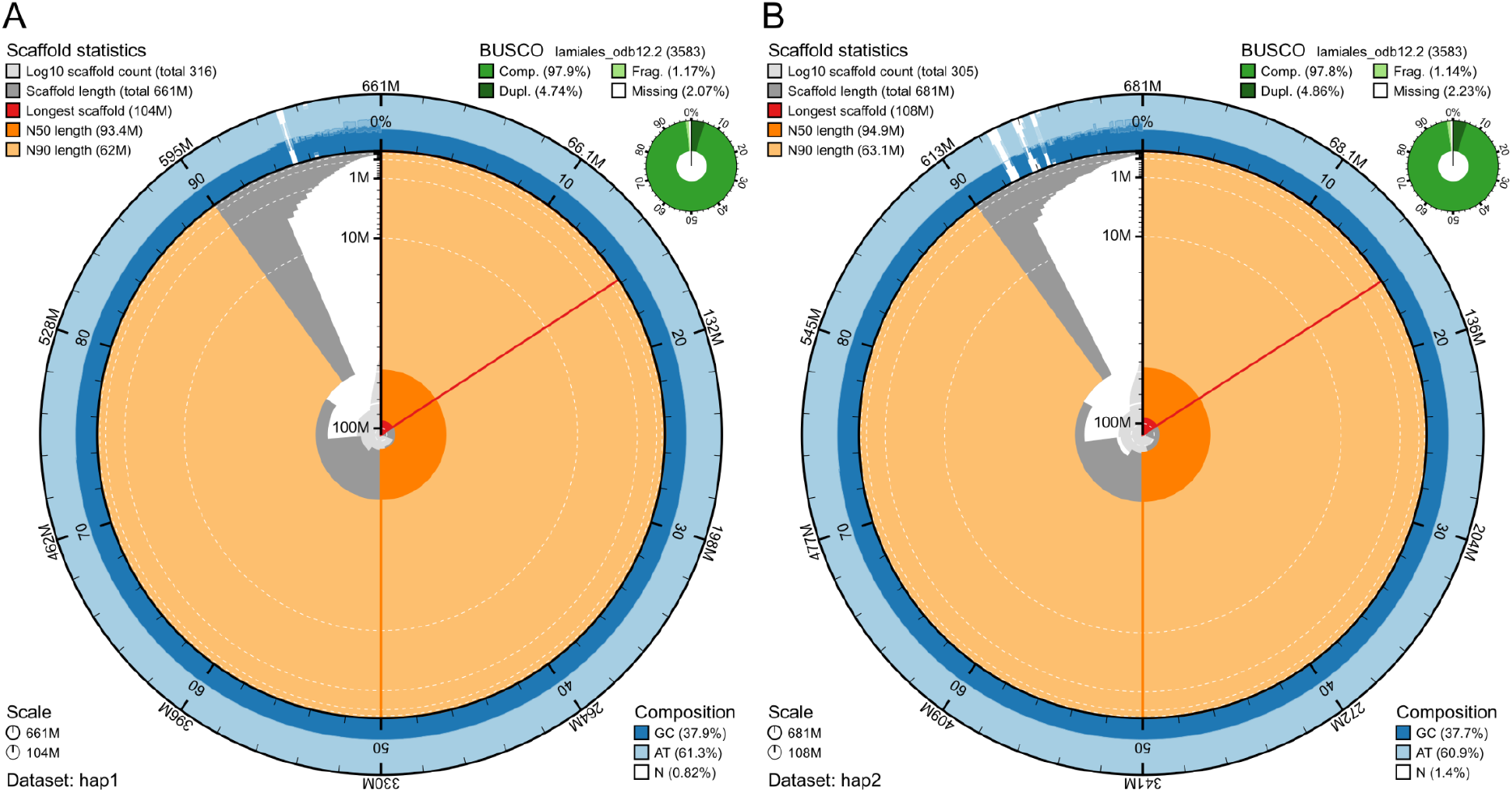
Metrics of the genome assemblies of *Dracocephalum ruyschiana* hap1 and hap2. A) Hap1. B) Hap2. The BlobToolKit Snailplots show N50 metrics and BUSCO gene completeness. The two outermost bands of the circle signify GC versus AT composition at 0.1% intervals, with mean, maximum, and minimum. The third outermost shows the N90 scaffold length, while the fourth is N50 scaffold length. The line from middle to second outermost band shows the size of the largest scaffold. All the scaffolds are arranged in a clockwise manner from largest to smallest, and shown in darker gray with white lines at different orders of magnitude, shown as a scale on the radius. The light gray shows the cumulative scaffold count. The scale inset in the lower left corner shows the total amount of sequence in the whole circle, and the fraction of the circle encompassed in the largest scaffold.

The Hi-C contact maps show well-resolved chromosome-scale scaffolds in both pseudo-haplotypes (Supplementary Figure 1).

When compared to a k-mer database of the Hi-C reads, hap1 had a k-mer completeness of 94.6%, hap2 of 94.5%, and combined they had a completeness of 95.6%. Further, hap1 had an assembly consensus quality value (QV) of 43.26 and hap2 of 43.62, where a QV of 40 corresponds to one error every 10,000 bp, or 99.99% accuracy compared to a k-mer database of the Hi-C reads (Table 2). Merqury k-mer spectra showed that most assembly k-mers were assigned to the expected shared or haplotype-specific categories, supporting the quality of the haplotype-resolved assembly (Supplementary Fig. 2). When comparing the two pseudo-haplotypes using minimap2, there are 1,267,805 nucleotide substitutions (0.28% of the aligned sequence), 128,736 deletions in hap2 compared to hap1 ranging from 1 bp to more than 1000 bp and 129,140 insertions from 1 bp to more than 1000 bp in size. A total of 38,141 and 38,617 protein-coding genes were annotated in hap1 and hap2, respectively (Table 2).

## Discussion

We present a chromosome-level, pseudo-haplotype-resolved genome assembly of northern dragonhead with seven chromosomes per pseudo-haplotype, consistent with the known karyotype (2n = 14). The assemblies show good contiguity, supported by their high scaffold N50 values, and high completeness, as indicated by BUSCO scores and combined k-mer completeness.

The sequenced individual exhibited relatively high heterozygosity (0.805%), highlighting the value of resolving both haplotypes to capture genomic variation within an individual. Comparison of the two pseudo-haplotypes revealed approximately 1.27 million nucleotide substitutions and numerous insertions and deletions within aligned regions. The chromosome-level assemblies and associated gene annotations provide a reference for future investigations of genetic diversity and differentiation across the species’ fragmented distribution, building on the regional genetic structure previously described by Kyrkjeeide et al. (2020).

*D. ruyschiana* is classified as Vulnerable in Norway (Kyrkjeeide et al., 2020; Solstad et al., 2021). Its conservation is also important for the endemic and endangered northern dragonhead sap beetle, *Thymogethes norvegicus*, which depends on the plant for reproduction and for which a complementary genome assembly has been generated (Strand et al., 2026). The beetle study also recovered two bacterial endosymbiont genomes, including *Symbiodolus*, whose predicted shikimate pathway to chorismate likely contributes to beetle nutrition. Together, these genomic resources provide a foundation for future research and conservation of *D. ruyschiana* and the specialized plant–beetle–microbe system it supports.

## Acknowledgements

This project received data management and infrastructure support from ELIXIR Norway, supported by the Research Council of Norway’s grants 322392 and 350529, the University of Bergen, the University of Oslo, the Arctic University of Norway in Tromsø, the Norwegian University of Science and Technology and the Norwegian University of Life Sciences: NMBU. The authors acknowledge support from the National Infrastructure for High Performance Computing and resources provided by Sigma2 as well as Data Storage in Norway (project NN8013K) for computational work. The Norwegian Sequencing Centre generated the sequencing data used in this project (http://sequencing.uio.no).

## Data Availability

Data generated for this study are available under ENA BioProject PRJEB90390. PacBio sequencing data for the northern dragonhead (ENA BioSample: SAMEA118426445) are deposited in ENA under ERR15095666, ERR15095667 and ERR15095668, while Illumina Hi-C sequencing data are deposited in ENA under ERR15365184. Pseudo-haplotype one can be found in ENA at PRJEB90104, while pseudo-haplotype two is PRJEB90389.

The genome assemblies and gene annotations are available at Zenodo: https://doi.org/10.5281/zenodo.23980571

