## Supplementary for "A chromosome-level genome assembly of northern dragonhead, *Dracocephalum ruyschiana* L., 1753"

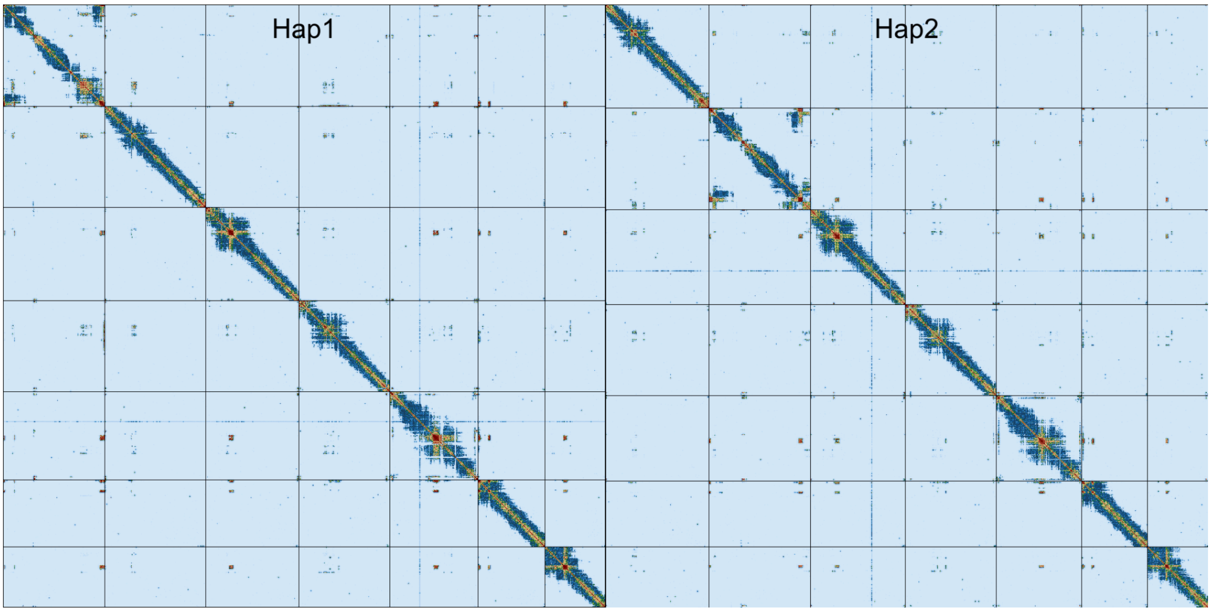


**Supplementary Figure 1: Hi-C contact maps for northern dragonhead, *Dracocephalum ruyschiana,* hap1 and hap2.** The contact map displays interaction frequencies between genomic regions, where darker shades represent a higher number of Hi-C contacts. The axes correspond to the coordinates along each assembly. The Hi-C contact maps were generated by mapping the Hi-C reads to the haplotype genomes using BWA-mem, generating a contact map using PretextMap and visualized using PretextSnapshot.

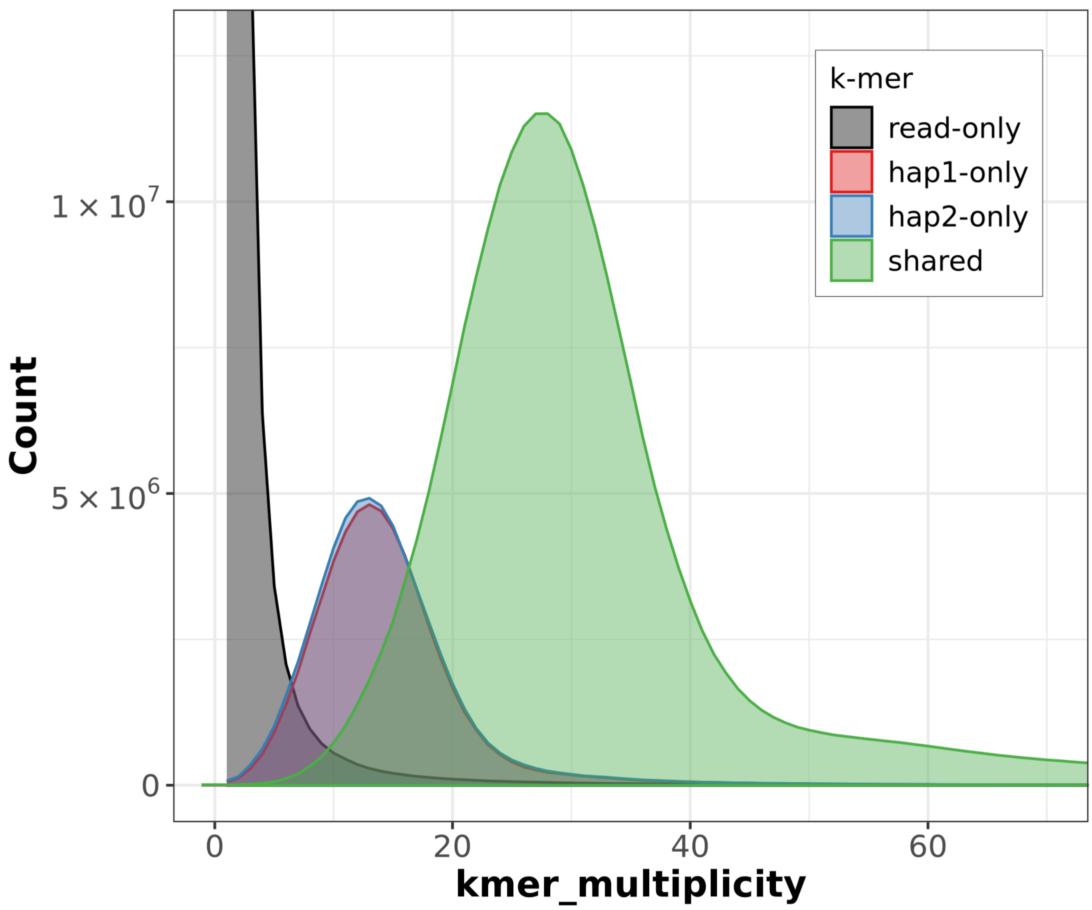


**Supplementary Figure 2: *K*-mer spectra of HiFi reads for northern dragonhead, *Dracocephalum ruyschiana*.** Distributions of *k*-mers found only in the reads (black), only in haplotype 1 (red), only in haplotype 2 (blue), or in both haplotypes (green). The y-axis represents the number of unique *k*-mers, while the x-axis represents the *k*-mer multiplicity (how often the *k*-mer is found in the set of HiFi reads).
